# Physiological caffeine concentrations do not affect enteric neural crest cell migration in a mouse model

**DOI:** 10.64898/2026.11.03.763904

**Authors:** Olivier Romito, Justine Peres, Monica Dicu, Emeline Lelong, Isabelle Le Parco, Nadège Bondurand, Nicolas R. Chevalier

## Abstract

The defect in enteric neural crest cell (ENCCs) migration that causes Hirschsprung disease (HD) in newborns is linked to genetic and likely also environmental factors. Among these, maternal coffee consumption has been associated with increased HD risk, yet the role of caffeine on ENCC migration has remained unexplored. Here, we show that although mouse ENCCs express caffeine-sensitive ADORA2A receptors, physiological doses of caffeine do not perturb endogenous ENCC calcium activity. We find that caffeine can modulate ENCC outgrowth in a 2D culture model but that it does not affect migration in a 3D collagen gel and in the ex-vivo embryonic gut. We conclude that physiological doses of caffeine do not hamper ENCC migration in the ex-vivo embryonic mouse intestine, thereby discarding a potential direct environmental mechanism of caffeine in HD.

## Introduction

The neural crest is a transient, highly migratory embryonic cell population that gives rise to a wide diversity of derivatives, including melanocytes, glia cells, and the enteric nervous system (Brandon et al., 2023; Bronner and LeDouarin, 2012; Cordero et al., 2011). Neural crest cells travel large distances within the embryo, delaminating and migrating collectively as streams of cells (Brandon et al., 2023; Bronner and LeDouarin, 2012; Cordero et al., 2011). Enteric neural crest cells (ENCCs) are responsible for building the enteric nervous system (ENS) (Bondurand et al., 2018; Laranjeira and Pachnis, 2009). In the mouse embryo, ENCCs enter the foregut at approximately embryonic day 9.5 (E9.5) and colonize the entire gastrointestinal tract by around E13.5 (Yu et al., 2024). This colonization process depends on continuous ENCC proliferation, migration, and responsiveness to biochemical cues, notably Glial cell-line derived neurotrophic factor (GDNF) and endothelin 3 (edn3), which are both produced and secreted by the surrounding gut mesenchyme (Morikawa et al., 2024; Young et al., 2001). We recently demonstrated that ENCC migration requires Ca^2+^ activity, which is triggered by edn3 and mediated by T-Type voltage-gated Ca^2+^ channels (Chevalier et al., 2026). Any disruption of these mechanisms can leave distal segments of the gut without ganglion cells, leading to Hirschsprung disease (HD) (Ardenne and Reitnauer, 1975; Bondurand et al., 2018; Mederer et al., 2020).

HD affects approximately 1 in 5000 live births (Morera and Nurko, 2024). The RET gene encodes the RET tyrosine kinase receptor of GDNF (Bondurand and Southard-Smith, 2016) and is the major gene involved in isolated HD. Mutations in GDNF/RET account for around 50% of familial cases and 15–20% of sporadic cases (Tomuschat and Puri, 2015). About half of HD cases do not however have a simple genetic explanation, pointing to more complex oligogenic inheritance and a likely contribution of gene-environment interactions during the time when ENCCs migrate in the developing gut. Among candidate environmental factors, maternal coffee consumption during pregnancy has been associated with an increased risk of HD in offspring with Down Syndrome: women who drank more than 3 cups of coffee per day showed a 6 time higher risk than lighter or non-consumers (Torfs and Christianson, 1999). Caffeine, the principal psychoactive compound in coffee (Burdan, 2015), is a modulator of cell migration in multiple cell types (Lee et al., 2026; Manica et al., 2024; Meisaprow et al., 2021) and is known to reach the fetal compartment during pregnancy (Rohweder et al., 2024). Whether caffeine can actually interfere with ENCC migration, and by which mechanism, has not previously been assessed.

Here, we investigated whether caffeine has the potential to act directly on ENCCs at physiological doses and whether this action translates into altered cell migration during gut colonization. We show that ENCCs express caffeine-sensitive ADORA2A adenosine receptors. In a 2D migration assay, caffeine significantly increased explant outgrowth and abolished GDNF-induced ENCC outgrowth. We did not however find any effect of caffeine on ENCC migration in 3D collagen gels or within the colonic mesenchyme. While caffeine could trigger Ca^2+^ activity in ENCCs migrating in 2D from explants, it did not affect ENCC Ca^2+^ activity in the native E11.5 gut. These findings indicate that caffeine can directly act on ENCCs and interfere with the GDNF pathway under specific (2D) experimental conditions, but that it is does not disrupt ENCC migration within the intact gut.

## Materials and methods

### Animals, dissection and ethics

We used the GCaMP6f NCC-specific mouse model previously described (Chevalier et al., 2026). Mice were bred and housed at the Institut Jacques Monod animal facility (Paris), with free access to food and water, controlled temperature (20–24°C), humidity (45–55%), and a 12h light/dark cycle (lights on 7am–7pm). Timed matings were set up in the evening and confirmed the following morning by vaginal plug detection. Pregnant dams were euthanized by cervical dislocation to collect embryos at E11.5, which were rapidly decapitated with surgical scissors. Procedures comply with CNRS/INSERM animal welfare guidelines. As embryo collection is a terminal, non-survival procedure, it does not require a specific ethics approval number. Guts were dissected from stomach to anus, with particular care to keep the mesentery intact. Samples were maintained in PBS with 1 mM Ca^2+^, 0.5 mM Mg^2+^, and 1% penicillin-streptomycin.

### Outgrowth assays

For the 2D outgrowth assay, E11.5 midgut segments (2-3 per gut) were cultured on 40 mm diameter culture-treated Petri dishes (TPP) and maintained for 48h at 37°C, 5% CO_2_ in 2 mL DMEM F12 supplemented with 10% FBS, with or without GDNF (10 ng/mL, Alomone Labs G-240), and/or caffeine (10 µM, Sigma C0750). For the 3D outgrowth assay, we first coated culture dishes with 0.5 mL of collagen (1 mg/mL, Cultrex) and let it gel for 20 min at 37°C. We then placed a single midgut explant in each dish and further covered it with 0.5 mL collagen, that was gelled for 45 min at 37°C, following the manufacturer’s protocol. This two-step procedure guaranteed that the explant was not in direct contact with the dish surface and that migration was thus completely 3D. 2 mL of DMEM F12 supplemented with 10% FBS and complemented with GDNF (± caffeine, as indicated) were added and the preparations cultured for 72h at 37°C, 5% CO_2_. The final GDNF concentration was always 10 ng/mL. For both 2D and 3D assays, the cell halo was revealed with a binocular (Leica) using a thin slit of transmitted light (pseudo-interferential contrast mode). Migration was quantified from the explant area (S2) and the total area including the cell halo (S1), yielding an average radial migration distance 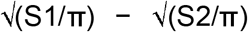, which corrects for halo asymmetry.

### Immunofluorescence

After 48h culture, 2D explants were fixed in 4% PFA/PBS (10 min), permeabilized/blocked in 0.1% Triton, 1% BSA/PBS (1h), and incubated with anti-SOX10 (mouse monoclonal, Sigma AMAB91297, 1:200) alongside with anti-ADORA2A (rabbit polyclonal, Abcam 3461, 1:100), 24h each, with 3 PBS washes between primary and secondary steps. Species-appropriate fluorophore-conjugated secondaries (Thermofisher) (1:500) were applied overnight before imaging.

### Data base analysis

Publicly available RNA-sequencing data from the Gene Expression Omnibus (GEO, accession GSE250526) were used to examine *Adora* receptor expression in ENCCs at E13.5 (Pai et al., 2023). Processed expression values (normalized RNA-seq dataset) were retrieved for *Adora1, Adora2a, Adora2b*, and *Adora3*, and compared across the four genes to identify the predominant *Adora* subtype expressed in ENCCs.

### Calcium imaging

We used a spinning-disk confocal setup (Olympus IX-81/Yokogawa CSU-X1, Andor ILE laser, Andor Zyla camera, 1392×1040 px) for calcium recordings. Embryonic guts were immobilized in low-melting point agarose. GCaMP6f fluorescence was excited at 488 nm and collected at 497–527 nm with a 10x objective, at 1 Hz, 300 ms exposure, for 5 min. Individual cells were investigated using ROI and mean gray value using ImageJ. Oscillations were quantified using CaSiAn software (CaSiAn · bio.tools).

### Whole-gut ENCC migration assay

Entire guts, from stomach to anus, were placed on 35 mm Petri dish pre-coated with 0.5 mL of 1% agarose (to prevent migration on the dish) and embedded in a second layer (0.5 mL) of agarose. Preparations were covered with 2 mL DMEM F12 supplemented with 10% FBS ± caffeine. Spinning-disk confocal z-stacks were acquired at culture start to locate the initial position of the ENCC wavefront, and after 48h, allowing precise measurement of the migration distance during the culture period.

### Statistics

All N values refer to independent embryos, and n refers to individual samples. Data were collected across ≥3 litters and ≥3 independent experimental days per condition, set according to expected effect variability. Statistical tests are specified in individual figure legends. Briefly, all Mann-Whitney were two-tailed and Tukey post-hoc were used for One-way Anova.

## Results

We first examined the expression of ADORA receptors, the primary targets of caffeine at physiological concentration (Han et al., 2020; Lemes Dos Santos Sanna et al., 2024). Analysis of publicly available RNA-seq data from the Gene Expression Omnibus (GSE250526) revealed that *Adora1, Adora2a*, and *Adora2b*, but not *Adora3* were expressed in ENCCs from E13.5 onward, with *Adora1* and *Adora2a* expressed at higher levels than *Adora2b* (Fig. 1A). Unfortunately, we did not find suitable data sets at earlier development stage. We confirmed ADORA2A expression by immunofluorescence at E11.5. We found that ADORA2A was distinctly expressed by GCaMP- and SOX10-positive ENCCs, but also by SOX10-negative mesenchymal cells (Fig. 1B).

**Figure 1:**
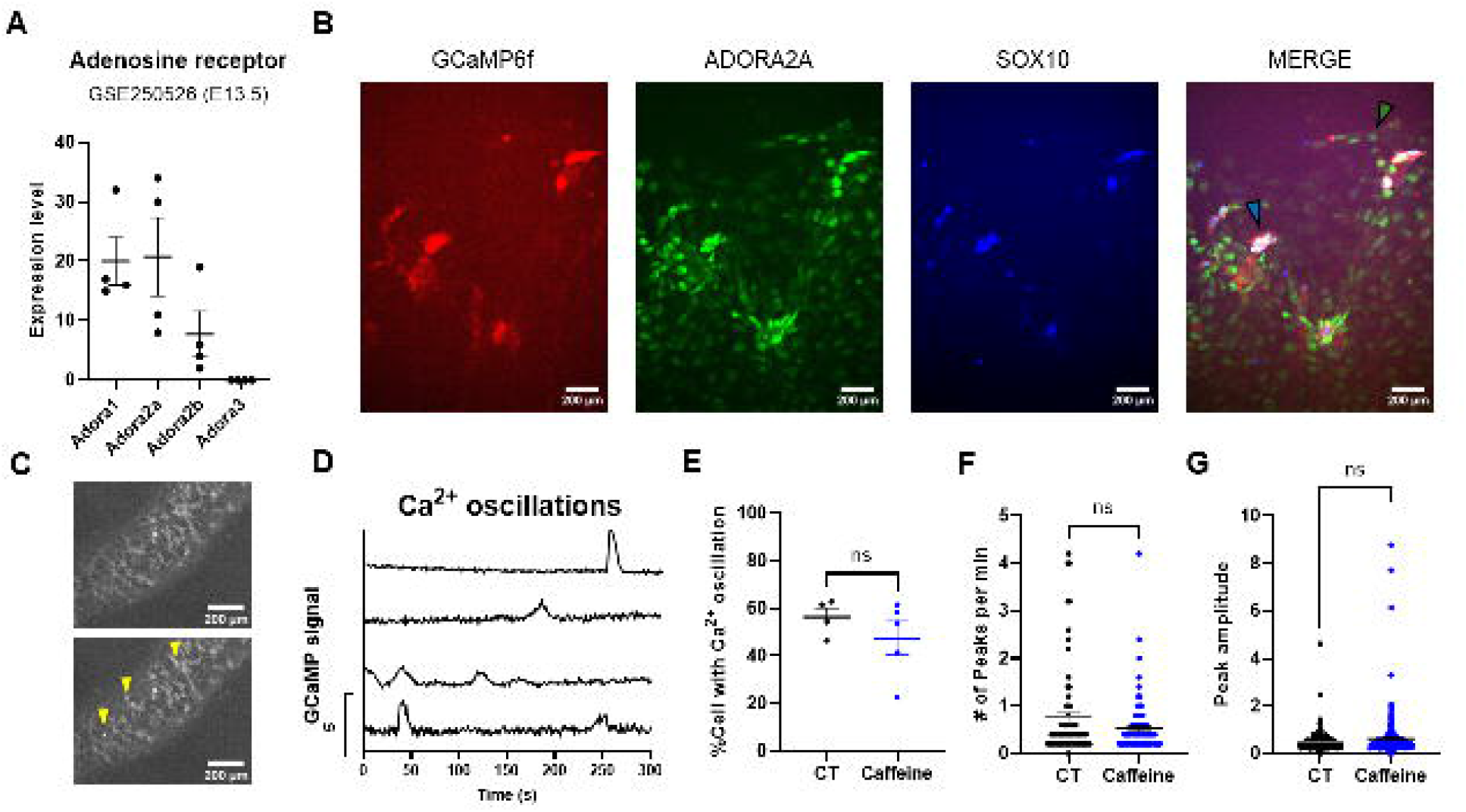
ENCCs express ADORA receptors during gut colonization window. A. mRNA analysis of *Adora1-3* at E13.5 using publicly available data base (GSE250526) (n=4). B. Immunofluorescence of GCaMP6f, ADORA2A and SOX10 from GCaMP6f NCC specific E11.5 embryonic mouse intestinal explants cultured for 48h in 2D (scale bar: 200 µm, N=3). C. GCaMP6f fluorescence in the E11.5 small intestine at 2 time points, yellow arrows indicate oscillatory cells (scale bar: 200 µm). D. Representative Ca^2+^ oscillations from individual cells in ENCC from GCaMP6f restricted non-treated E11.5 mouse embryo. (E-G) Proportion of ENCCs displaying Ca^2+^ oscillations between control (N=4 guts) and 48h caffeine (10 µM, N=5 guts) treated whole intestines (Mann-Whitney: p =0.3175). F. Frequency of Ca^2+^ oscillations in ENCC from entire intestine in control (N=4 guts, n=70 cells) or caffeine (N=5 guts, n=105 cells) treated samples for 48h (Mann-Whitney: p=0.5249). G. Peak amplitude in ENCC from entire intestine in control (N=4 guts, n=267 cells) or caffeine (N=5 guts, n=271 cells) treated samples for 48h (Mann-Whitney: p=0.9962). Data are represented as mean ± SEM.

Caffeine has been reported to modulate Ca^2+^ signaling in smooth muscle cells (SMCs) through an ADORA-dependent mechanism (Alexander and Cheung, 1994). As we recently showed that ENCC migration relies on intracellular pathways comparable to those of smooth muscle cells (Chevalier et al., 2026), we investigated the effect of caffeine on Ca^2+^ oscillations. In 3D explant culture of E11.5 segments cultured for 2 days, we found that caffeine could trigger a transient increase of Ca^2+^ in ENCCs at 10 µM (n=2/6) or at 100 µM (n=1/6) (Video S1). This confirms that ADORA receptors on ENCCs are functional and receptive to caffeine. This effect did not however occur in native E11.5 guts, since acute administration of 10 µM caffeine did not modify the proportion of cells exhibiting Ca^2+^ activity, or the average frequency and amplitude of oscillations (Fig. 1C-G).

We next asked whether caffeine could affect ENCC migration, first resorting to 2D explant culture (Fig. 2). Under all conditions, including untreated controls, cells were detected outside the explant, consistent with a baseline level of outgrowth sustained by growth factors present in the culture medium (Fig. 2A,B). In 2D explant culture, the migration halo consists of both mesenchymal cells and ENCCs: we confirmed this by fluorescence imaging and also by the fact that not-yet-colonized E11.5 colon segments could give rise to a migration halo exclusively of mesenchymal origin (Fig. 2C). Working on small intestine segments and in the absence of GDNF, we found that caffeine (10 µM) significantly increased baseline outgrowth compared to controls (Fig. 2A and 2B). As expected, GDNF increased outgrowth, with a stronger effect than caffeine (Fig. 2A,B). Surprisingly, in the presence of GDNF, caffeine reduced the GDNF-induced outgrowth (Fig. 2A,B). When the same experiment was performed with colon explants, i.e., with a cell halo that was exclusively mesenchymal, caffeine did not alter halo size (Fig.2C,D). This indicates that the detrimental effects of caffeine on GDNF-induced outgrowth targeted the ENCCs, and not the surrounding mesenchyme.

**Figure 2:**
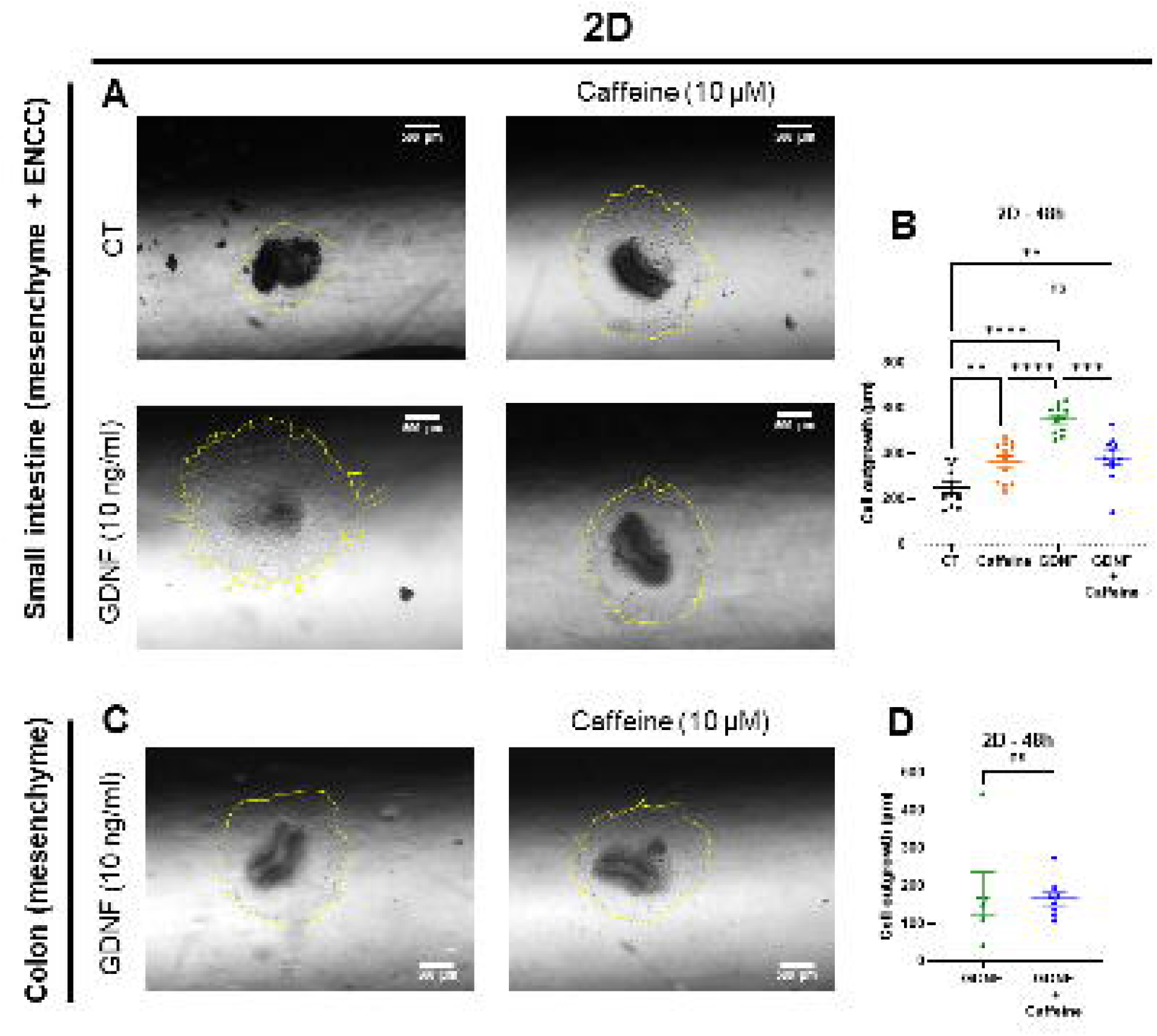
Caffeine inhibits GDNF-induced 2D gut explant outgrowth in an ENCC-dependent manner. A. Representative photographs of small intestine segment from E11.5 embryos, cultured in 2D for 48h, with or without caffeine (10 µM), and/or GDNF (10 ng/ml) (scale bar: 500 µm). B. Average outgrowth distance from the small intestine explant after 48h of culture (11<N<12 guts, One-way Anova: p<0,0001, F= 25.12). C. Representative photographs of colon segments from E11.5 embryos, cultured for in 2D for 48h with GDNF (10 ng/ml), with or without caffeine (10 µM) (scale bar: 500 µm). D. Average outgrowth distance from the colon explants after 48h of culture (6<N<8 guts, Mann-Whitney: p = 0,6620). Data are represented as mean ± SEM.

Because these first experiments hinted at a potential interference of caffeine with the GDNF-RET pathway, we next assessed migration in a more physiological 3D collagen gel assay. Only ENCCs penetrate the collagen lattice, removing the mesenchymal contribution. We found that caffeine (10 µM and 100 µM) did not affect GDNF-induced 3D outgrowth, compared to GDNF treated samples (Fig. 3A,B). We finally examined ENCC migration after 48h across the cecum and colon of E11.5 ex-vivo whole guts. In these conditions, caffeine may influence both the ENCCs and the mesenchymal compartment within which they migrate. We found that the average migration distance was ∼650 µm after 48h (∼15 µm/h), and was similar for control and caffeine treated (10 µM) samples (Fig. 3C, 3D).

**Figure 3:**
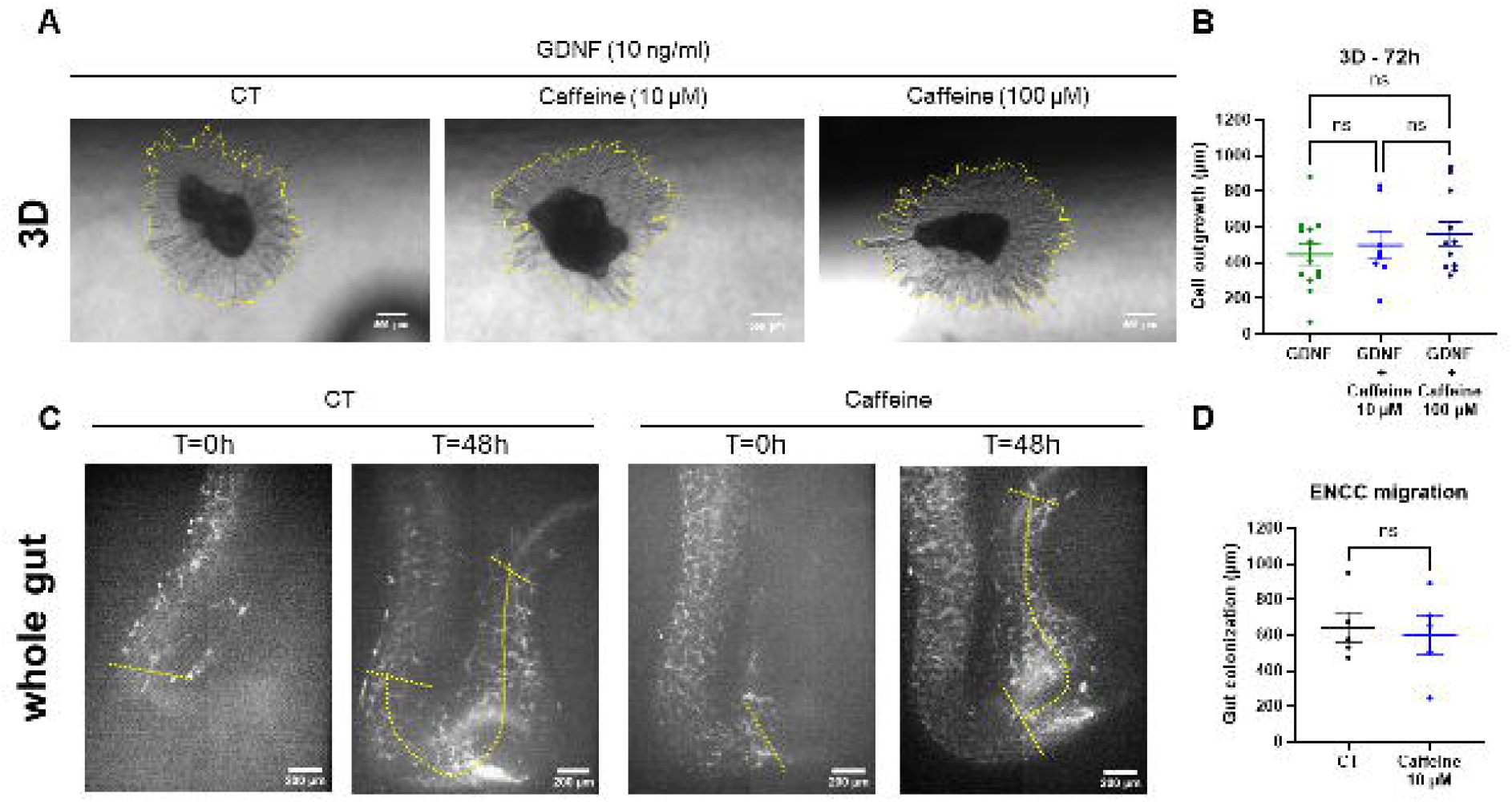
Caffeine does not affect ENCC migration in 3D collagen gels or in the whole ex-vivo intestine. A. Representative photographs of E11.5 small intestine segments cultured for 72h in a 3D collagen gel with GDNF (10 ng/ml), and treated with or without caffeine (10 µM; 100 µM) (scale bar: 500 µm). B. Average outgrowth distance from the explant after 72h of culture (8<N<13 guts, One-way Anova: p= 0,4396, F= 0,8456). C. Representative projected z-stacks of whole-guts at T=0h and after 48h culture, with or without caffeine (10 µM) (scale bar: 200 µm). D. Average ENCC gut colonization distance (CT: N=5 guts, Caffeine: N=6 guts, Mann-Whitney: p >0,9999). Data are represented as mean ± SEM.

### Discussion and conclusion

Caffeine is widely consumed in the diet of Western populations (Reyes and Cornelis, 2018). It is present in coffee, tea, cocoa, as well as in cola and energy, chocolate, confectionery and some medications drinks (Wikoff et al., 2017). Coffee and tea are major sources of caffeine intake (Landais et al., 2018), and caffeine exposure is frequent during pregnancy. Most women continue to drink coffee and tea before they become aware of their pregnancy, and usually decrease their intake thereafter (Rohweder et al., 2024). This is particularly relevant to ENS development because human ENCCs enter the developing foregut at gestational week 4 and progressively colonize the entire gut by week 7-8 (Wallace and Burns, 2005). Thus, ENCC migration happens at a period of pregnancy when the mother may not have yet modified her dietary habits. Because caffeine crosses the placenta and maternal caffeine clearance is reduced during pregnancy (Struniewicz et al., 2025), current recommendations aim to limit maternal caffeine intake to approximately 200 mg/day (2-3 cups). This sets a biologically relevant concentration range for examining potential alterations to embryo development (Struniewicz et al., 2025). A clinical report associated maternal consumption of more than 3 cups of coffee per day during pregnancy with an increased risk of Hirschsprung disease (HD) in offspring with Down syndrome (Torfs and Christianson, 1999). On a total of 687 infants with Down Syndrome, 1.5% percent exhibited HD, ∼10 times more than the average disease prevalence in the general population. This number increased to 6.9% when the mothers consumed 4 cups of coffee par day during the periconceptional period (Torfs and Christianson, 1999). Although this association does not establish causality, it raises the possibility that caffeine, the principal psychoactive component of coffee, may directly interfere with developmental processes required for enteric nervous system (ENS) formation.

In this study we used caffeine at 10 µM (∼maximum serum concentration after ingestion of 1 cup (Martínez-López et al., 2014)) and 100 µM. At these doses, caffeine acts predominantly as a non-selective antagonist of adenosine receptors (Ec50 ∼10 µM (McLellan et al., 2016)), rather than through phosphodiesterase inhibition (Ec50 ∼400 µM (McLellan et al., 2016)) or other lower-affinity targets such as ryanodine receptors (Ec50 ≥1 mM (McLellan et al., 2016)). We showed that ENCCs and the surrounding mesenchymal compartment express adenosine ADORA2A receptors during the period of gut colonization. We recently reported that ENCC migration is intimately linked to their Ca^2+^ activity (Chevalier et al., 2026) and since caffeine is known to alter Ca^2+^ in a variety of cells (Endo, 1977; McPhersonx et al., 1991; Missiaen et al., 1994; Parker and Ivorra, 1991), we examined its effect on migrating ENCCs. We found that physiological caffeine concentration (10 µM) can promote a transient Ca^2+^ surge in 3D explants, similar to its effect in smooth muscle cells (Guerrero et al., 1994; Porta et al., 2011). This effect shows that adenosine receptors on ENCCs may be stimulated by caffeine, likely by blocking endogenous adenosine binding to ADORA receptors. However, caffeine did not alter Ca^2+^ activity in the entire E11.5 mouse gut.

We further found that caffeine can affect ENCC migration differently depending on whether migration occurs on a 2D substrate or in a 3D collagen gel. While the 2D assay revealed a possible interference of caffeine with GDNF signaling, more physiologic collagen gel and whole-mount 3D assays did not show any impact of caffeine on ENCC migration. This discrepancy may plainly be due to the fact that 2D migration involves interspersed and interacting mesenchymal and ENC cells, whereas the 3D and whole-mount assay involve only ENCCs. It also underlines that migration mechanisms are fundamentally different in these two configurations (Chevalier et al., 2016): 3D migration happens by the formation of chains of cells and by the degradation of the ECM by metalloprotease, while cell-substrate adhesions play a more prominent role in 2D. The ability of caffeine to increase outgrowth in 2D explants is consistent with previous observations showing that caffeine can increase cell migration in vascular cells (Lee et al., 2026; Wang et al., 2021). Specifically, caffeine (10 µM to 100 µM) increases endothelial cell migration in a cAMP/PKA dependent manner (Wang et al., 2021), which may constitute the mechanism observed in our study.

We showed that although ENCCs can be receptive to caffeine via adenosine receptors, physiological concentrations of caffeine are not detrimental to their 3D migration inside collagen gels or within the embryonic mouse cecum and colon. These experiments do not exclude the possibility that caffeine could have effects on earlier stages of NCC migration. Also, we have only investigated the effect of caffeine. Coffee contains numerous other bioactive compounds, including chlorogenic acids, trigonelline, and diterpenes, which could, in isolation, or synergistically with caffeine, promote an effect on cell migration (Iwamoto et al., 2019; Liao et al., 2015; Makino et al., 2021). Finally, ex-vivo experiments can substantially differ from in vivo. The vasodilating effect of caffeine is for example mediated via activation of sympathetic nerves at the central nervous system level, and not via a direct effect of caffeine on smooth muscle (Echeverri et al., 2010; Higgins and Babu, 2013). Future work based on chronic in vivo coffee or caffeine administration will be needed to determine whether, and how, maternal coffee consumption can translate into a risk factor for HD.

## Data availability

The dataset supporting the conclusions of this article is included within the article and its additional file.

## Authors contribution

OR wrote the manuscript that was critically reviewed by NB and NC. OR and JP performed the experiments. MD, EL and ILP were in charge of the animal work. NC supervised the project.

## Funding and acknowledgements

This study was supported by grants from the University of Paris Cité, France (Electrocre).

